# Accurate characterization of expanded tandem repeat length and sequence through whole genome long-read sequencing on PromethION

**DOI:** 10.1101/439026

**Authors:** Arne De Roeck, Wouter De Coster, Liene Bossaerts, Rita Cacace, Tim De Pooter, Jasper Van Dongen, Svenn D’Hert, Peter De Rijk, Mojca Strazisar, Christine Van Broeckhoven, Kristel Sleegers

## Abstract

Tandem repeats (TRs) can cause disease through their length, sequence motif interruptions, and nucleotide modifications. For many TRs, however, these features are very difficult - if not impossible - to assess, requiring low-throughput and labor-intensive assays. One example is a VNTR in *ABCA7* for which we recently discovered that expanded alleles strongly increase risk of Alzheimer’s disease. Here, we investigated the potential of long-read whole genome sequencing to surmount these challenges, using the high-throughput PromethION platform from Oxford Nanopore Technologies. To overcome the limitations of conventional base calling and alignment, we developed an algorithm to study the TR size and sequence directly on raw PromethION current data.

We report the long-read sequencing of multiple human genomes (n = 11) using only a single sequencing run and flow cell per individual. With the use of fresh DNA extractions, DNA shearing to approximately 20kb and size selection, we obtained an average output of 70 gigabases (Gb) per flow cell, corresponding to a 21x genome coverage, and a maximum yield of 98 Gb (30x genome coverage). All *ABCA7* VNTR alleles, including expansions up to 10,000 bases, were spanned by long sequencing reads, validated by Southern blotting. Classical approaches of TR length estimation suffered from low accuracy, low precision, DNA strand effects and/or inability to call pathogenic repeat expansions. In contrast, our novel NanoSatellite algorithm, which circumvents base calling by using dynamic time warping on raw PromethION current data, achieved more than 90% accuracy and high precision (5.6% relative standard deviation) of TR length estimation, and detected all clinically relevant repeat expansions. In addition, we identified alternative TR sequence motifs with high consistency, allowing determination of TR sequence and distinction of VNTR alleles with homozygous length.

In conclusion, we validated the robustness of single-experiment whole genome long-read sequencing on PromethION, a prerequisite for application of long-read sequencing in the clinic. In addition, we outperformed Southern blotting, enabling improved characterization of the role of expanded *ABCA7* VNTR alleles in Alzheimer’s disease, and opening new opportunities for TR research.

## Introduction

Half of the human genome is estimated to consist of repetitive DNA elements. These are categorized as interspersed repeats (e.g. *Alu, LINE* elements and segmental duplications) and tandem repeats (TRs). The latter includes short tandem repeats (STRs; a.k.a. microsatellites) which have 1-6bp motifs and variable number of tandem repeats (VNTRs; a.k.a. minisatellites) with repeat unit length > 6bp. Compared to non-repetitive DNA, our knowledge on repeats is lagging behind severely; mostly due to technological limitations^1^.

Currently, we know of approximately 50 TRs that affect disease, primarily neurological. STRs are particularly involved in rare diseases with high penetrance due to repeat expansions, and VNTRs are mostly associated with common complex disorders^2,3^. Most often, pathological and benign alleles are categorized based on an arbitrary repeat length cutoff^3^. In reality, however, the disease associated effects of TRs are more complex. Firstly, one TR can be involved in multiple diseases^2,3^. Secondly, increasing length of expanded repeats can lead to increased severity of the phenotype or anticipation^3^. Thirdly, repeats can be interrupted by alternative sequence motifs, which influences repeat stability and modifies associated phenotypes^4–8^. Lastly, CpG dinucleotides in TRs can be methylated which may contribute to disease development^9–13^.

A comprehensive analysis of TRs therefore requires accurate length estimation, nucleotide sequence determination and preferably analysis of epigenetic modifications. Unfortunately, the techniques most often used to study TRs in clinical diagnosis and basic research, i.e. Southern blotting and repeat-primed PCR, only provide an estimation of length, with accuracy inversely correlated with repeat size. In addition, these methods only target one TR locus at a time, and have high turnaround times. Currently, there is no assay to study tandem repeats simultaneously, let alone on a human genome scale^14^.

Next generation sequencing reads from the most conventional platforms (e.g. Illumina) are too short to directly resolve TRs. In the last two years, new algorithms were developed to circumvent this limitation, which enables screening for potential STR expansions. However, these length estimations lack accuracy and validation with gold standard techniques like Southern blotting is still necessary^14^.

A solution can theoretically be obtained with long-read sequencing. These technologies have rapidly improved in the last years with both Pacific Biosciences (PacBio) and Oxford Nanopore Technologies (ONT) now providing the opportunity to perform human whole genome long-read sequencing. Long sequencing reads can span entire (expanded) TR alleles, providing TR length, nucleotide composition, and the possibility to detect nucleotide modifications. In practice, however, the higher sequencing error rates of long-read sequencing may limit this application and necessitates evaluation^15^. ONT sequencing has several characteristics which makes it particularly attractive to study TRs. It is based on direct sensing of nucleotides and does not require DNA polymerization. As such, ONT sequencing suffers less from a GC coverage bias in the often GC-rich TRs^16^, and nucleotide modifications can be directly detected. Secondly, there is no technical maximum on read length and the sequencing quality does not decay with increasing length^17^. Thirdly, real-time data processing on ONT devices can lower the turnaround time^18^. Lastly, with the release of the high-throughput PromethION sequencing platform in 2018, ONT now provides the most cost effective human whole genome long-read sequencing option.

A few reports exist on the use of long-read sequencing in tandem repeats. However, these are limited to small-scale tests on plasmids, specifically amplified genomic regions, and/or do not include expanded TR alleles^8,19,20^. Only one study attempted whole genome sequencing of an individual with a *C9orf72* repeat expansion; ONT sequencing (on the low throughput MinION device) yielded 3x coverage without reads spanning the expanded allele^21^.

The aim of this study was twofold. First, we tested the feasibility and robustness of long-read human genome sequencing on the recently released high-throughput PromethION sequencing device (ONT) by sequencing DNA from eleven individuals. Second, we evaluated the use of this data to characterize TR length, sequence, and nucleotide modifications. We focused our analyses on an *ABCA7* VNTR, for which we recently discovered that expanded alleles are a strong risk factor for Alzheimer’s disease^22^. This VNTR (chr19:1049437-1050028, hg19) has a high GC-content, a 25bp repeat unit with frequent nucleotide substitutions and insertions (consensus motif embedded in Figure 1), and the total repeat size can reach more than 10,000 bp. Prior to this study only Southern blotting could be used to approximate the length of this challenging TR.

**Figure 1:**
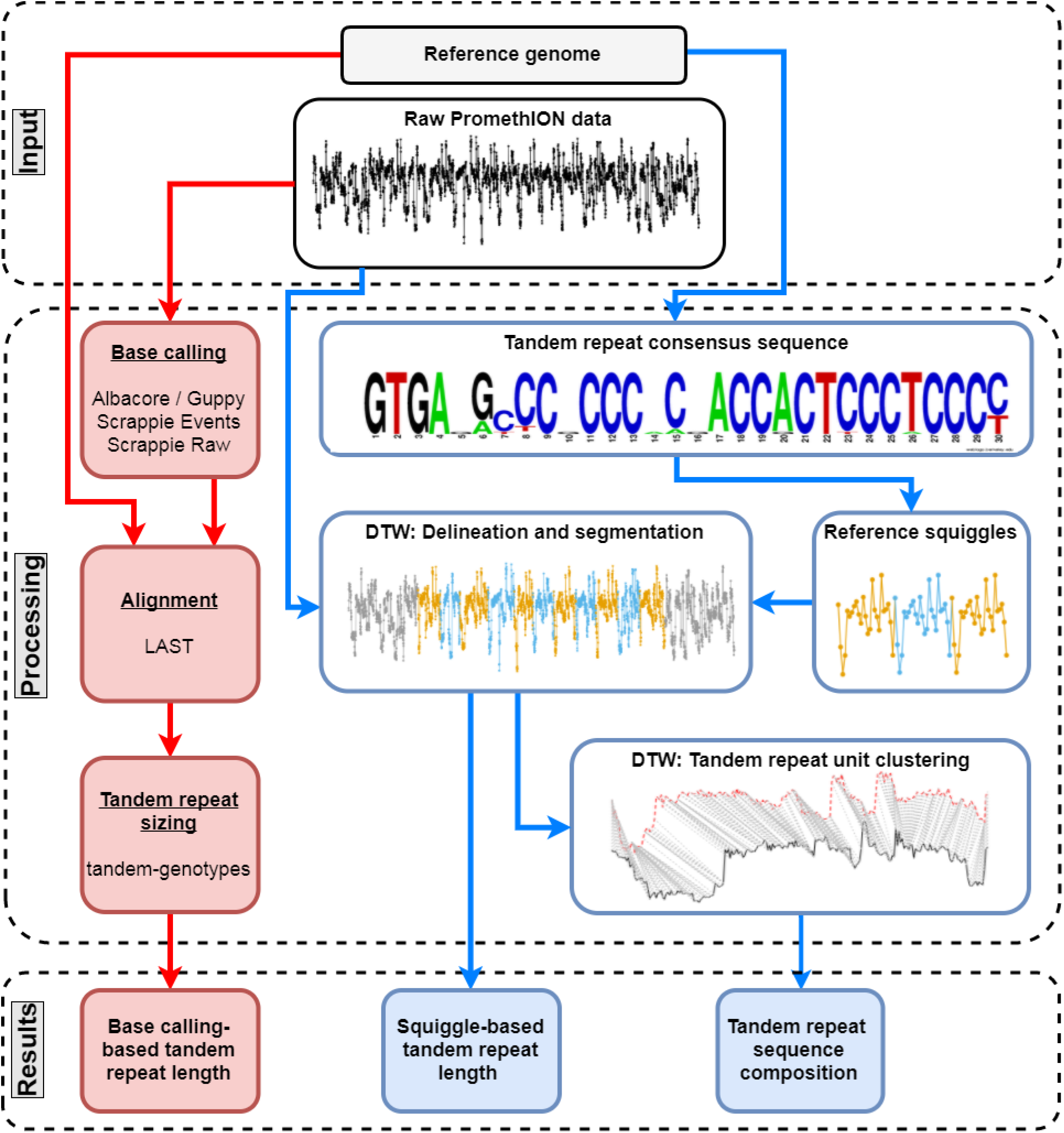
Tandem repeat analysis methods. To extract TR length and sequence information from PromethION data, we used a base calling (red) and our squiggle-based NanoSatellite (blue) approach. Consecutive steps are shown in bold with below the names of the used bioinformatics tools or squiggle illustrations. The “Raw PromethION data” illustration corresponds to a partial PromethION squiggle from a single read spanning the *ABCA7* VNTR. The “Tandem repeat consensus sequence” figure was obtained from *De Roeck et al*., 2018. The height of each nucleotide corresponds to its frequency on that position^22^. In the “Reference Squiggles” figure, three *ABCA7* VNTR units (alternating colors) are shown based on Scrappie current estimation. After DTW with raw PromethION data and reference squiggles, the TR is delineated from the flanking sequence and segmented into individual TR units (alternating colors in the “Delineation and segmentation” figure). The final “Tandem repeat unit clustering” figure depicts the DTW process between two TR units. Each current measurement from a TR unit (red or black) is matched (gray lines) to a current measurement from the other TR unit.

## Materials and methods

### Study population

We performed long-read whole genome sequencing on DNA from eleven individuals (Table 1), using the PromethION sequencing platform (Oxford Nanopore Technologies (ONT), Oxford, United Kingdom). Ten individuals (Subject01 till Subject10) were recruited in the context of the Belgian Neurology (BELNEU) Consortium^22,23^ and consisted of AD patients (n = 6), an FTLD patient, a family member at risk of developing dementia, and healthy elderly control individuals (n = 2). All participants and/or their legal guardian provided written informed consent for participation in genetic studies. The study protocols were approved by the ethics committees of the Antwerp University Hospital and the participating neurological centers at the different hospitals of the BELNEU consortium and by the University of Antwerp. For all individuals, Epstein-Barr virus transformed lymphoblastoid cell lines (LCLs) were available. In addition, we included the previously described NA19240 PromethION sequencing dataset^24^. *ABCA7* VNTR lengths were determined in all individuals with Southern blotting as previously described^22^.

**Table 1:** Summary of individuals included in this study. DNA was either extracted with QIAmp (Qiagen) or on a Magtration robotic platform (PSS). An alpha (α) and beta (β) PromethION device were used. Two chemistries were applied: SQK-LSK108 (108) or SQK-LSK109 (109). Yield and mean read length were calculated with NanoPack^41^ after Albacore or Guppy base calling. kb = kilobases, Gb = gigabases, N50 = 50% of the total sequencing dataset is contained in reads equal or larger than this value, a = DNA was not sheared for this individual, b = An eight year old DNA extraction was used, c = SQK-LSK109 was used for 4 flow cells and for 1 flow cell was run with SQK-LSK108. d = 6 debug PromethION flow cells and 7 MinION flow cells were used.

| Individual | ABCA7<br>VNTR<br>lengths (kb) |  | DNA source | PromethION<br>device | Sequencing<br>chemistry | Number<br>of flow<br>cells | Yield<br>(Gb) | Read length<br>N50 (kb) |
| --- | --- | --- | --- | --- | --- | --- | --- | --- |
| Subject01 | 5.1 | 4.5 | QIAmp | $\alpha$ | 108 | 6+7<sup>d</sup> | 74.4 | 7.0 |
| Subject02 | 1.8 | 1.8 | QIAmp | $\alpha$ | 108 | 1 | 61.3 | 11.2 |
| Subject03 | 4.7 | 2.3 | QIAmp | $\alpha$ | 109 | 1 | 72.5 | 16.3 |
| Subject04 | 9.7 | 0.8 | QIAmp | $\alpha$ | 109 | 1 | 85.5 | 11.5 |
| Subject05 | 10.7 | 1.7 | QIAmp | $\alpha$ | 109 | 1 | 88.5 | 11.9 |
| Subject06 | 2.3 | 0.3 | QIAmp | $\alpha$ | 109 | 1 | 98.0 | 16.2 |
| Subject07 | 3.3 | 3.3 | Magtration | $\alpha$ | 109 | 1 | 77.7 | 10.6 |
| Subject08 | 2.3 | 0.4 | Magtration | $\beta$ | 109 | 1 | 56.2 | 14.7 |
| Subject09<sup>a</sup> | 5.5 | 4.3 | Magtration | $\beta$ | 109 | 1 | 43.2 | 29.3 |
| Subject10<sup>b</sup> | 8.7 | 2.8 | Magtration | $\beta$ | 109 | 1 | 48.9 | 9.1 |
| NA19240 | 4.5 | 2.2 | Magtration | $\beta$ | 109<sup>c</sup> | 5 | 220.0 | 15.8 |

### DNA preparation for PromethION sequencing

LCL were cultured with 1640 RPMI medium (Thermo Fisher Scientific), supplemented with 15% Fetal Bovine Serum, 2mM L-Glutamine, 1mM Sodium Pyruvate, 100 IU/mL Penicilline, and Streptavidine. Cell counting was performed on a Luna II (Logos Biosystems, South Korea) automated cell counter. Five million cells per individual were then resuspended in PBS. To establish the optimal protocol for DNA preparation for PromethION sequencing, different procedures for DNA extraction and fragmentation were tested. Extraction of DNA was done according to the manufacturer’s protocol with either QIAmp DNA Blood mini spin columns (Qiagen, Hilden, Germany) or automated extraction on a Magtration 8LX platform (Precision System Science (PSS), Matsudo, Japan) (Table 1). RNA degradation was carried out with RNase A during cell lysis in the QIAmp extraction protocol, and after completion of the Magtration protocol.

The extracted DNA was fragmented to a mean length of 15kb or 20kb with Megaruptor (Diagenode, Liège, Belgium), with the exception of Subject09 and three aliquots of NA19240 which remained unsheared. All DNA was size selected on BluePippin (Sage Science, Beverly, USA), with a high pass protocol retaining all fragments above a minimum size cutoff, which ranged from 7 to 10kb. DNA was subsequently purified with Agencourt AMPure XP beads (Beckman Coulter, Brea, USA) in a 1:1 volume ratio. DNA size analysis was conducted on a Fragment Analyzer with DNF-464 High Sensitivity Large fragment 50kb kit, as specified by the manufacturer (Agilent Technologies, Santa Clara, USA).

### ONT library preparation and sequencing

Different sequencing set-ups were applied as outlined in Table 1 due to frequent improvements of ONT sequencing chemistry, flow cells and PromethION sequencing devices.

We followed the “1D Genomic DNA by Ligation sequencing on PromethION” protocol (Oxford Nanopore Technologies) according to the latest sequencing kit version (SQK-LSK108 or SQK-LSK109), with slightly increased incubation times during end-preparation, purification and final elution to increase yield. Briefly, DNA was first repaired and dA-tailed with NEBNext FFPE DNA Repair Mix (New England Biolabs (NEB), Ipswich, USA) and NEBNext End repair / dA-tailing (NEB). This was either performed serially (SQK-LSK108), or in a single step (SQK-LSK109). Purification was done with AMPure XP beads. Subsequently, ONT adapters were ligated to the DNA library with the NEB Blunt/TA Ligase Master Mix (SQK-LSK108), or with the NEBNext Quick Ligation Module and the ONT supplied “Ligation buffer” (SQK-LSK109). AMPure XP beads were then used for clean-up together with ONT’s “Adapter Bead Binding Buffer” (SQK-LSK108) or ONT’s “L(ong) Fragment Buffer” (SQK-LSK109). DNA was eluted in the supplied Elution Buffer and loaded on all four inlets of a primed PromethION flow cell.

The flow cells (FLO-PRO001), were either part of an initial debug phase for early PromethION optimization, or commercially available flow cells. For Subject01, six debug flow cells were used, while the other samples were sequenced on a single commercial flow cell, with the exception of NA19240 for which the goal was to obtain very high genome coverage depth by using multiple flow cells (Table1). Upon arrival, all flow cells were subjected to quality control and were used for sequencing within a week. With the exception of debug flow cells, all flow cells used for sequencing, had at least 6000 available pores.

During the course of this study, sequencing was conducted on two PromethION devices: an alpha and a beta unit. The main difference between both was the incorporated GPU computational module in the beta device, which allowed real-time base calling.

During the PromethION optimization phase, we also sequenced DNA from Subject01 on seven MinION flow cells (Table 1) to compare performance of the DNA extractions and library preparations on both platforms. The “1D Genomic DNA by Ligation sequencing on MinION” protocol was followed using SQK-LSK108 chemistry, R9.4 flow cells (FLO-MIN106) and a Mk1 MinION platform (MIN-101B).

### Data analysis

ONT flow cells contain protein nanopores connected to an application-specific integrated circuit (ASIC). As the DNA passes through the pore, nucleotide sequences are shifted, resulting in changes of ionic current that are detected by the ASIC. Each sequenced DNA fragment is therefore represented by a series of current levels, also known as a “squiggle” (illustration embedded in Figure 1). Conventionally, these squiggles are then base called with the use of neuronal network based software^25^. Albacore (ONT) was the first base caller available to the public and is currently most widely used. Recently, ONT released Guppy, which is similar to Albacore, but optimized for faster computation on GPU and embedded in beta PromethION sequencing devices. Lastly, ONT also provides the developmental base caller Scrappie (https://github.com/nanoporetech/scrappie), which has two modes of base calling: “events” and “raw”.

All PromethION sequencing reads were first processed using conventional base calling and genome alignment procedures: data from the alpha and beta PromethION devices were respectively base called with Albacore v2.2.5 (ONT) and Guppy v1.4.0 (ONT). Subsequent alignment was performed with minimap2^26^ with hg19 (GRCh37) as reference genome. NanoPack was used to summarize experiment metrics^27^.

#### ABCA7 VNTR analysis using existing methods for TR sizing

For purpose of comparison of existing methods for TR length determination, reads aligning to a 100 kb genome region containing the *ABCA7* VNTR and flanking sequences (chr19:1000000-1100000) were rebase called with Albacore (v2.2.5), Scrappie events (v1.3.1-f31cada), and Scrappie raw (v1.3.1-f31cada) (Figure 1). Alignment was then carried out on each of the three base called datasets using LAST v941^28^ with base caller-specific trained LAST parameters. Calculation of repeat length was performed with tandem-genotypes^20^. Tandem-genotypes estimates the number of tandem repeat units as the difference in length between sequencing reads and the reference sequence, divided by the consensus repeat unit size. To determine absolute repeat units per sequencing read, we added the number of tandem repeat units in the reference (23.2 for the *ABCA7* VNTR) defined by Tandem Repeats Finder (TRF)^29^. Computation was parallelized using gnu parallel^30^.

We evaluated the ability of sequencing reads processed by these three base callers to resolve TR sequence composition. All reads spanning the *ABCA7* VNTR (according to tandem-genotypes analysis) were processed with the TRF algorithm^31^ using lenient parameters to account for base calling errors: a matching weight of 2, mismatching penalty of 3, indel penalty of 5, match probability of 80, indel probability of 10, and minimum alignment score of 14. To determine the *ABCA7* VNTR pattern per sequencing read, we considered repeats with a pattern size between 18 and 32bp and selected the repeat with the highest copy number. Next, per base caller and DNA strand, we counted the number of reads with an *ABCA7* VNTR pattern, and calculated average sequence composition metrics.

### Squiggle based *ABCA7* VNTR data analysis

Existing methods for TR length determination as described above depend on base calling and alignment accuracy; both of which are often suboptimal in low-complexity and/or repetitive sequences. To overcome these challenges, we designed “NanoSatellite”, a novel pattern recognition algorithm, which bypasses base calling and alignment, and performs direct TR analysis on raw PromethION squiggles (Figure 1). Most of this method is based on consecutive rounds of Dynamic Time Warping (DTW), a dynamic programming algorithm to find the optimal alignment between two (unevenly spaced) time series. DTW is used in many different applications, such as speech recognition, analysis of electrocardiograms, and it is also the main constituent of “Read Until”, an algorithm designed to enrich sequences of interest on an Oxford Nanopore sequencing platform by quickly identifying patterns at the squiggle level^32^.

#### Code and data availability

All code is freely available on GitHub (https://github.com/arnederoeck/NanoSatellite). All *ABCA7* VNTR spanning raw current PromethION fast5 files are publicly available via study PRJEB29458 on the European Nucleotide Archive (https://www.ebi.ac.uk/ena/data/view/PRJEB29458). Data accession to NA19240 sequencing data is reported in De Coster et al., 2018^24^.

#### Generation of reference squiggles

To enable pattern recognition on raw PromethION data, we first translated DNA nucleotide sequences to an estimated squiggle pattern. We used TRF to delineate TRs of interest in the genome (chr19:1049437-1050028 for the *ABCA7* VNTR). Next, we extracted 250 bp of flanking sequence on both sides of the TR. We converted the consensus TR motif defined by TRF (GTGAGCCCCCCACCACTCCCTCCCC for the *ABCA7* VNTR), as well as the flanking sequences and their respective reverse complements to approximate squiggles using the “squiggle” module of Scrappie (v1.3.1-f31cada).

#### Tandem repeat delineation

Using the reference squiggles, TR length determination was performed on raw PromethION data with DTW by first finemapping the TR boundaries, and subsequently segmenting the TR into individual TR units. First, we selected TR spanning reads for which tandem-genotypes was able to estimate tandem repeat lengths and we included reads aligning on both sides of the TR after minimap2 alignment. Raw squiggle data was extracted from the original fast5 files using the rhdf5 package^33^ in R^34^. The extracted current levels were split in overlapping windows. These windows and the reference squiggles were then z-scale normalized. DTW of strand-matched flanking sequence reference squiggles and windowed sequencing reads was performed using the dtw R package^35^ with both sides unfixed. Since each nucleotide is represented by one data point in reference squiggles and multiple current measurements in raw PromethION data, we applied the Minimum Variance Matching (MVM) algorithm, with an MVM step pattern of 25 (each reference squiggle data point is allowed to match 25 current measurements or less)^36^. We selected the windows with the lowest DTW distance and fine mapped the TR start and end with DTW using a reference squiggle composed of flanking sequence and multiple TR units.

#### Tandem repeat segmentation

Subsequently, the delineated PromethION TR squiggle was split up in individual TR units using a strand-matched reference squiggle composed of five TR units. DTW with a 25 MVM step pattern was started at both ends of the TR with one end fixed and an open end towards the center of the TR. In both sets, three TR units were defined, after which the process was repeated with the fixed end matching the last defined unit of the previous cycle, until both sets crossed each other. Overlapping segmentations were resolved by choosing the segmentation with the lowest DTW distance. The average distance from TR segmentation in all sequencing reads was recorded. Sequencing reads with an average distance larger than 1.5 times the interquartile range from the 75^th^ percentile were discarded.

#### Tandem repeat unit clustering and sequence determination

For the purpose of identification of the underlying nucleotide sequence, all TR unit squiggles were clustered using the dtwclust package^37^ in R^38^. Strand-specific distance matrices were calculated with symmetric DTW distances. Subsequently, TR units were clustered into two groups using hierarchical clustering with Ward’s method. Centroids for each cluster were extracted using the partition around medoids (PAM) method. To determine the corresponding TR unit sequence, we compared the centroids to reference squiggles of known alternative VNTR motifs based on TRF analysis of the *ABCA7* VNTR in the reference genome. We reconstructed the sequencing reads as a chain of TR unit clusters and created a consensus sequence through alignment with the msa package^39^.

### Tandem repeat statistics and visualization

We evaluated TR length estimation of the three tested base callers combined with tandem-genotypes and the squiggle approach by calculating several metrics for each. All TR length estimations by PromethION sequencing reads (*r_a,i_,) were assigned per method and per subject to a TR allele *a*, with *i* ranging from 1 to the total number of reads per allele (*n_a_*). The average length of all reads per allele is denoted as *r*̅_a_* with standard deviation *s_a_*, and the Southern blotting length per allele as *L_a_*. The accuracy (the degree to which PromethION length estimations correspond to Southern blotting length estimation) per allele corresponds to 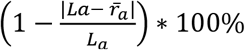 and a method-specific accuracy was calculated by taking the median of all allele accuracies. As a measure for precision (corresponding to the closeness of estimates from individual reads originating from the same allele) we determined the relative standard deviation per allele as 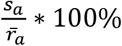 and calculated the method-specific median. Alleles with less than 2 reads in one of the four analysis methods were not included in these accuracy and precision calculations. All visualizations were made with ggplot2^40^ in R^38^.

## Results

### Human long-read genome sequencing with a single flow cell

We sequenced native genomic DNA from eleven individuals on an Oxford Nanopore PromethION sequencing platform. On average, we attained 70,195,516,005 bases (70.2 Gb; ~ 22x genome coverage) output per PromethION flow cell, with a maximum of 98.0 Gb (30.6x) (Table 1). The lowest yields were obtained for Subject09 (43.2 Gb) and Subject10 (48.9 Gb). For these individuals we respectively used unsheared DNA, or an eight year old DNA extraction instead of mechanically fragmented and recently extracted DNA as used in the other sequencing libraries. The final yield of each sequencing run was influenced by the number of available nanopores over time (Figure S1).

Overall, the mean read length N50 was 14.0 kb (half of the total number of bases originates from reads with a read length larger than or equal to the N50 value). Sequence length distributions differed between sequencing datasets (Figure S2). With the exception of Subject10 - whose old DNA extraction resulted in shorter reads - read lengths correlated strongly (R^2^ = 0.96) with the DNA fragment size prior to ONT library preparation and sequencing (Figure S3). Apart from Subject01 which we sequenced during the PromethION optimization phase, all sequencing datasets had a similar distribution of quality and an overall median identity to the reference genome of 86% (Figure S4).

### Base calling-based tandem repeat length determination

We evaluated PromethION analysis methods on their ability to resolve different *ABCA7* VNTR lengths varying from 300 bases (~ 12 repeat units) to more than 10,000 bases (~ 400 repeat units), as previously determined by Southern blotting (Table 1). On the one hand, we tested the performance of existing tandem repeat analysis methods for which we evaluated three base callers combined with the *tandem-genotypes* algorithm (Figure 1). We observed that length estimates based on *Albacore* - the most commonly used base caller - were underestimated with the largest deviation observed for VNTR spanning reads originating from the guanine-rich negative DNA strand (Figure 2a), resulting in low accuracy and precision (Table 2). Using the *Scrappie events* base caller, we observed better TR length estimation accuracy and a smaller, but persistent, strand bias effect (Figure 2a, Figure S5, Table 2). Overall, length estimations by *Scrappie raw* most closely approached the expected Southern blotting lengths with the lowest relative standard deviation, and the highest number of identified VNTR spanning reads (Figure 2a, Figure S5, Table 2). Nevertheless, while *Scrappie raw* showed the best results overall, it performed worst for the expanded *ABCA7* VNTRs (> ~229 repeat units). *Scrappie raw* was unable to identify spanning reads for the largest allele of Subject04 (Figure S5d) and only one spanning read was found for both Subject05 (Figure 2b) and Subject10 (Figure S5h), which had strongly deviating length estimates. Changes to base calling models and parameters in *Scrappie raw* did not result in improvements (personal communication with Tim Massingham, ONT).

**Figure 2:**
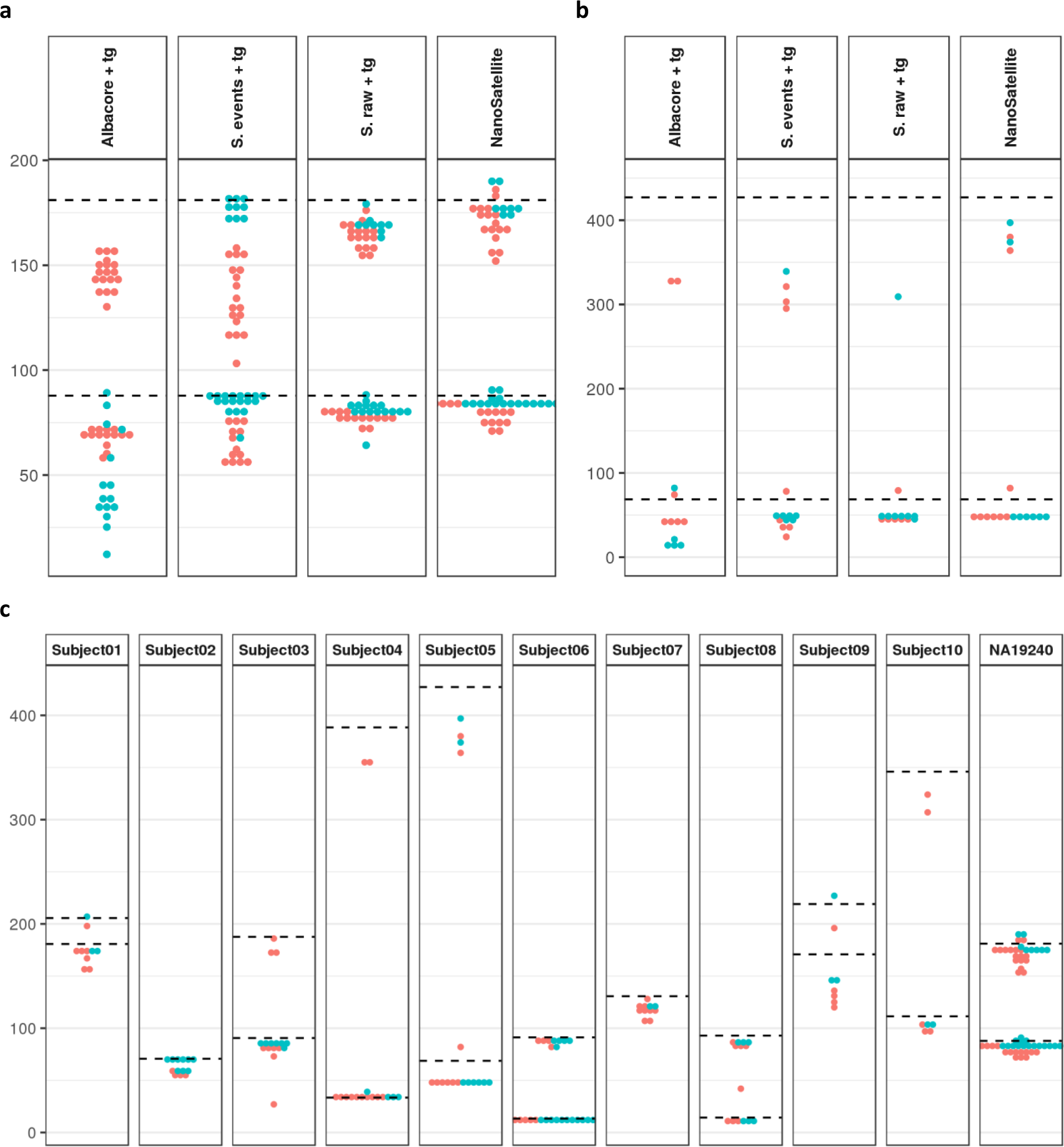
*ABCA7* VNTR length estimates. TR length estimates (the number of tandem repeat units is depicted on the y-axis) per positive strand (red) or negative strand (blue) PromethION sequencing reads (dots) are shown in comparison to the Southern blotting lengths (dashed lines). **(a)** Comparison of the four methods (Albacore + tandem-genotypes (tg), Scrappie (S.) events + tg, S. raw, and NanoSatellite) for NA19240, the individual with most sequencing reads. **(b)** Comparison of the four methods for Subject05, the individual with the largest expanded *ABCA7* VNTR allele. **(c)** NanoSatellite *ABCA7* VNTR length estimations for all individuals.

**Table 2:** Evaluation of tandem repeat analysis methods on the *ABCA7* VNTR. Accuracy corresponds to the degree of resemblance of the average length estimation and Southern blotting length. The relative standard deviation depicts the spread of length estimates to the mean. The total number of *ABCA7* VNTR spanning reads detected per method is shown under “Number of spanning reads”. In “Expanded read detection”, the detection rate of all known expanded reads is shown, with 100% corresponding to detection of all expanded reads.

| Method | Accuracy (%) | Relative standard deviation (%) | Number of spanning reads | Expanded read detection (%) | Sequence composition |
| --- | --- | --- | --- | --- | --- |
| Albacore + tandem-genotypes | 65.6 | 36.0 | 154 | 63 | low consistency |
| Scrappie events + tandem-genotypes | 83.3 | 15.5 | 177 | 88 | low consistency |
| Scrappie raw + tandem-genotypes | 87.7 | **5.5** | 181 | 25 | low consistency |
| NanoSatellite | **90.3** | **5.6** | **194** | **100** | **high consistency** |

### Novel squiggle-based algorithm to improve tandem repeat length determination

To circumvent errors introduced by base calling and downstream alignment processing steps, we developed “NanoSatellite”, a novel algorithm resolving tandem repeats directly on raw PromethION squiggle data, using DTW (Figure 1). Tandem repeat squiggles were reliably distinguished from the flanking sequence and were subsequently segmented in tandem repeat units. Using this approach, we identified more VNTR spanning sequencing reads than the conventional methods described above, and we were able to resolve all VNTR alleles in each sequencing dataset (Figure 2c). Read length estimations were more accurate compared to conventional methods and precision was similar to *Scrappie raw*. In addition, we observed consistent results across all VNTR lengths and DNA strands, with the highest detection rate of expanded VNTR alleles (Figure 2c, Table 2).

### Consistent tandem repeat sequence determination with squiggle clustering

To detect *ABCA7* VNTR units with an alternative sequencing motif, we first analyzed the nucleotide sequences generated by conventional tandem repeat analysis methods. The tandem repeat was detected in most spanning sequencing reads produced by all three base callers. However, in general, the consensus size was smaller than expected, and substantially more mismatches and indels were observed (Table S1), which makes reliable TR sequence determination infeasible. *Albacore* in addition produced a VNTR sequence with a strongly diverging sequence composition (Table S1).

We assessed whether alternative sequence motifs can be differentiated on the squiggle level by NanoSatellite. We used unsupervised hierarchical clustering to classify the squiggle tandem repeat units. While more alternative sequences may be present, we separated in two clusters per DNA strand, which could be clearly differentiated from each other (Figure S6 and Figure S7). Since multiple nucleotides contribute to the current measurement at a given time (i.e. approximately 5 nucleotides for the nanopores used in R9.4 flow cells), a nucleotide change can have different effects on current measurements based on the surrounding nucleotide composition. A base substitution or indel can therefore result in different effect sizes in the positive or negative DNA strand direction. Clustering of positive squiggles was mainly driven by a guanine insertion, or cytosine to adenine substitution at nucleotide ten of the consensus motif (Figure 3a and Figure 3c), whereas negative strand clustering was most strongly affected by a cytosine to thymidine substitution at nucleotide 21 (corresponding to the fourth nucleotide in the positive strand) (Figure 3b and Figure 3d).

**Figure 3:**
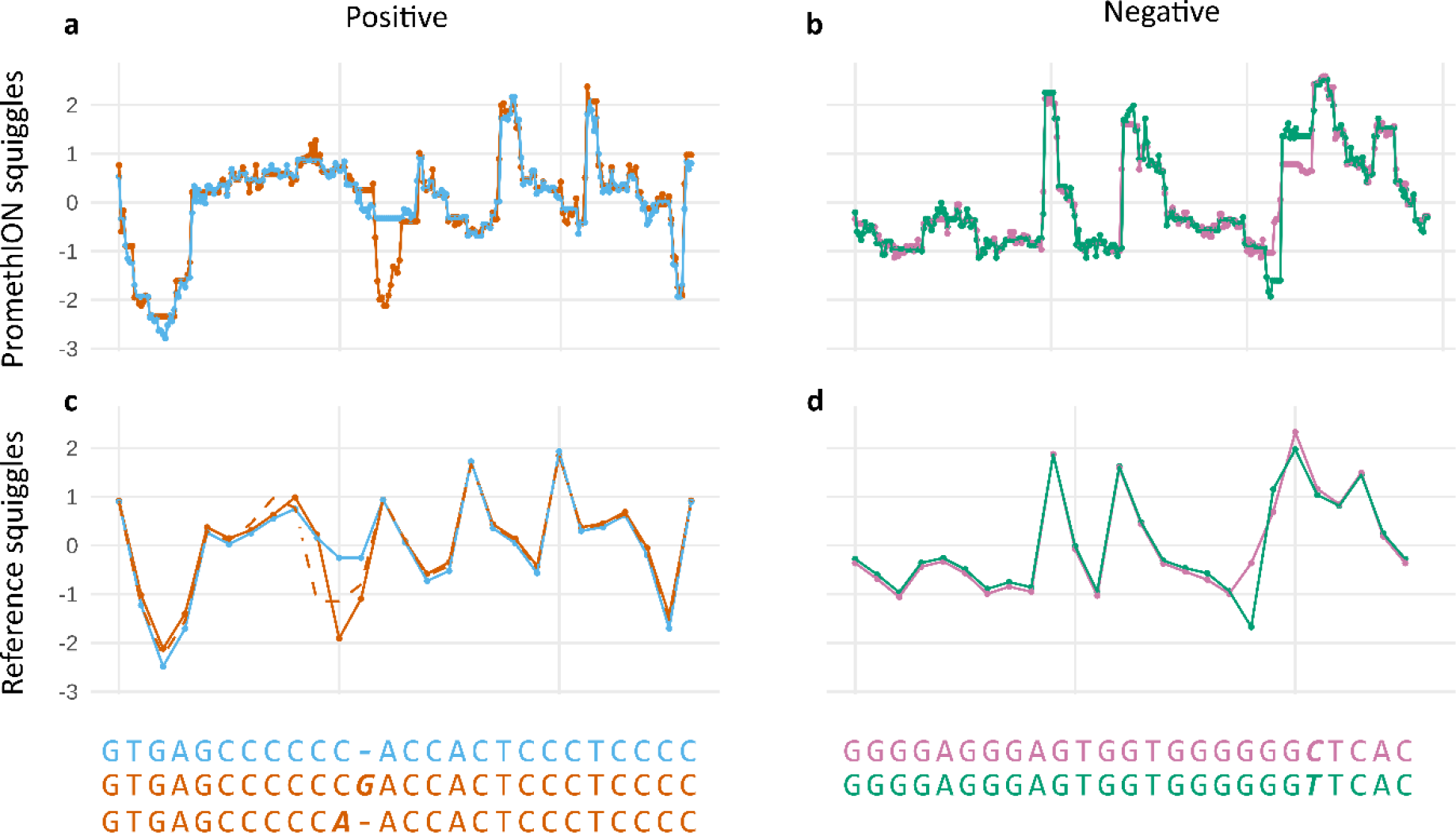
*ABCA7* VNTR squiggle clustering by NanoSatellite. Centroids are shown which were extracted from hierarchical *ABCA7* VNTR squiggle unit clusters originating from positive **(a)** or negative **(b)** DNA strands. Each cluster is shown in a different color. We compared these centroids to positive **(c)** and negative **(d)** reference squiggles with corresponding sequence motifs shown below. The top sequences (blue and purple) correspond to the expected VNTR motifs, while the sequences below (orange and green) contain nucleotide differences (in bold and italic). The alternative (orange) cluster observed in panel **a**, contains two alternative alleles: a guanine insertion (solid orange line in panel **c**) and a cytosine to adenine substitution (dashed orange line in panel **c**).

### Alternative sequence motifs can be used for fine-typing VNTRs

We determined the sequence of the original PromethION reads by ordering the TR unit clusters. We observed a high consistency in clustering patterns between different sequencing reads originating from the same VNTR allele (Figure 4a). Based on Southern blotting, only one VNTR length was observed for Subject02 and Subject07, compatible with homozygosity. In line with this, squiggle-based length estimation only identified reads corresponding to the Southern blot-based VNTR length (Figure 2c), however, by examining the read clustering patterns, we could distinguish two alleles that were indeed close in length, but had a different sequence composition (Figure 4b). While expanded alleles have fewer spanning sequencing reads due to their long lengths, we were able to determine a consistent consensus pattern, which differed between individuals (Figure 4c).

**Figure 4:**
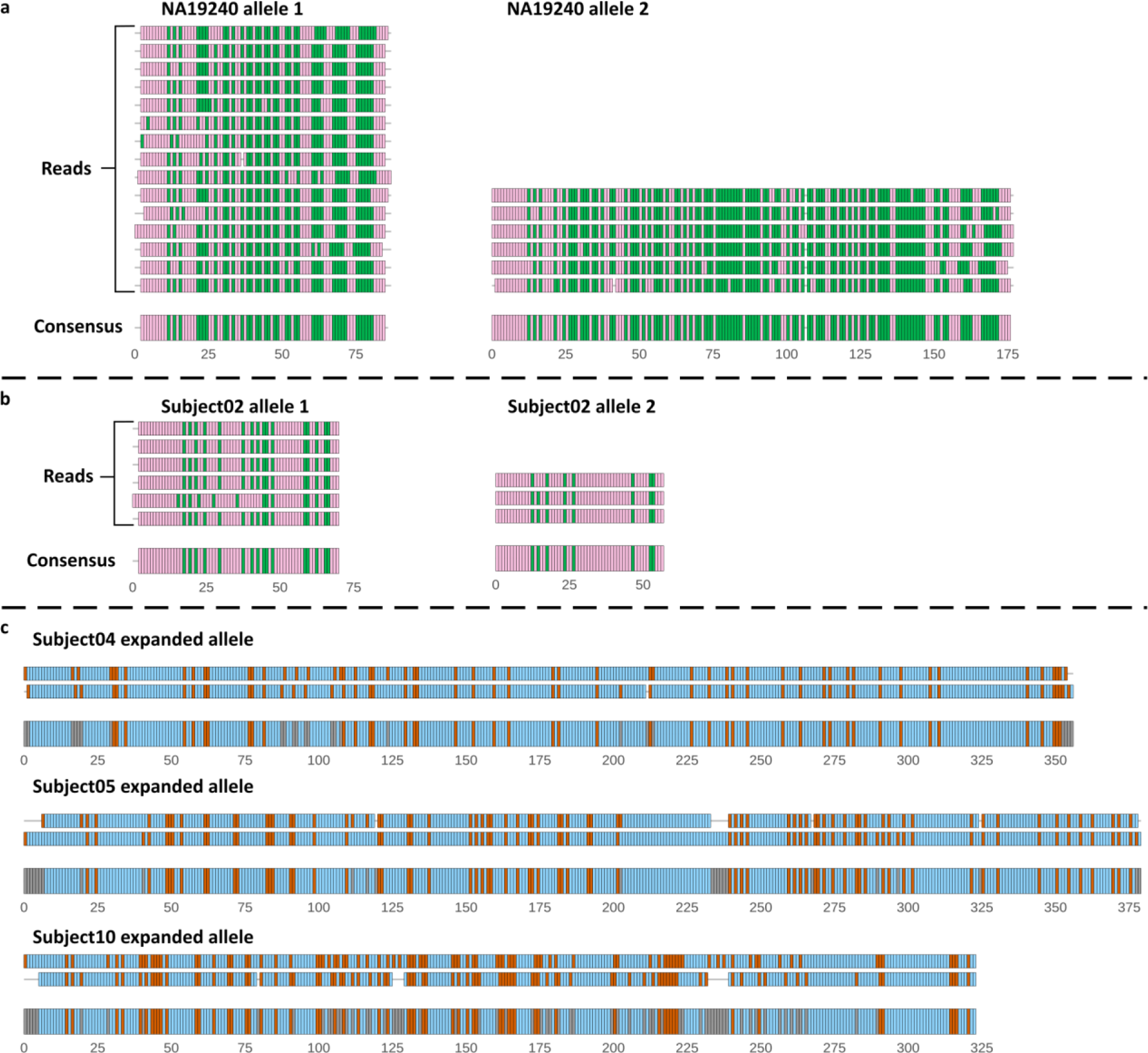
*ABCA7* VNTR sequence reconstruction based on squiggle clusters. Alignments are shown with individual PromethION sequencing reads (narrow segments) and a consensus (broad segments), as annotated in panels (a) and (b). Each rectangle corresponds to a tandem repeat unit. Colors correspond to the tandem repeat unit cluster as assigned in Figure 3. **(a)** Negative reads originating from the two NA19240 alleles. For both alleles of NA19240, two reads with deviating TR length were removed for clearer visualization. **(b)** Negative reads corresponding to both alleles of Subject02, for whom a single VNTR length is observed in Southern blotting. **(c)** Positive reads corresponding to the expanded alleles of Subject04, Subject05, and Subject10

## Discussion

*ABCA7* VNTR expansions result in a > 4-fold increased risk of Alzheimer’s disease^22^, yet the technological challenges of investigating TR sequences have so far precluded further research and large-scale screening. Long-read sequencing has the potential to overcome these issues on the condition that sufficient yield and read lengths are consistently obtained to traverse tandem repeats, and algorithms exist that can accurately size even the largest alleles. For the first time, we demonstrate the feasibility of long-read sequencing of multiple human genomes with a single sequencing run per individual; achieving up to 98 Gb output per PromethION flow cell. Furthermore, we developed an algorithm to study tandem repeats on a raw squiggle level and observed improved determination of read length and sequence composition compared to existing algorithms, which lacked accuracy, precision and/or ability to call large expanded alleles. For all datasets - even those with a relatively low yield, or relatively low mean read length due to DNA fragmentation - we were able to accurately determine repeat length for both *ABCA7* VNTR alleles, which ranged from 300bp to more than 10,000bp. The robust performance and high yield of single PromethION sequencing run combined with the high accuracy and precision of our novel algorithm suggest that future high-throughput detection and screening of TRs such as the *ABCA7* VNTR is attainable, both for research and clinical purposes.

### PromethION sequencing

With an average of 70.2 Gb per PromethION flow cell, we obtained substantially higher yields than the most recently published long-read human genomes. Generation of a 91.2 Gb genome on MinION (ONT) required 39 flow cells (2.3 Gb per flow cell on average)^42^, and 225 SMRTcells were required to produce a 236 Gb genome on a PacBio RSII instrument (1.0 Gb per SMRTcell)^43^. While we show that a single sequencing run is sufficient for long-read human genome sequencing, this study highlights several technical aspects that influence final yield. First, we observed the lowest yield for the sequencing run with the longest DNA fragments, for which we did not mechanically shear the DNA. For the same amount of DNA, longer fragments result in fewer available free ends to make contact to the nanopores and initiate sequencing. While we corrected for this by raising the amount of DNA loaded on a flow cell, this in turn could lead to suboptimal yield due to an increase in viscosity. In addition, the available free ends could be masked from the pore by steric hindrance of long DNA molecules. Hence we opted for shearing of DNA in most DNA libraries. The ideal shearing length to obtain satisfactory read lengths and high yields is not yet established and project dependent. We observed no clear relation between N50 read length and yield (Table 1); hence shearing at 20 kb (our highest shearing length) on a Megaruptor (Diagenode) was most optimal in our experience.

Second, the amount of DNA loaded on a flow cell makes a difference. Too few molecules result in suboptimal occupation of “open” pores, and too much DNA with attached motor proteins can cause faster depletion of ATP in the sequencing buffer. We adhered to suggested loading amounts as much as possible, yet these recommendations have changed over time. Third, the number of good sequencing pores at the start of sequencing varied due to manufacturing, shipment, and/or storage conditions. Nevertheless all sequencing runs were performed on flow cells with at least 6000 good pores at the start. Fourth, the rate of pore decline varied, which in turn is determined by several parameters. Interfering chemicals introduced during DNA extraction or library preparation can affect the stability of the flow cell (e.g. detergents can disrupt the membrane in which the nanopores are embedded), pores can collapse, and pores can get blocked. Several efforts by ONT have addressed these issues while this study was in progress including removal of detergents from consumables, brief current reversals during sequencing to unblock pores, and dynamic re-evaluation of “good” sequencing pores instead of fixed 16 hour time intervals, as used during the PromethION sequencing runs described in this study. Therefore, we anticipate yield will increase even further in future experiments.

We performed PromethION sequencing using DNA extracted by different methods, sequencing chemistries, and sequencing devices. We observed resembling read quality and similar resolution of *ABCA7* VNTR assessment for all datasets, demonstrating a robust generation of raw sequencing data by the platform. The read length correlated well to DNA fragment size when sequencing fresh DNA extractions. When we used an eight year old DNA extraction, however, shorter read lengths were observed. A potential explanation is the introduction of single strand nicks due to DNA degradation. Library preparation and DNA length determination are not affected by these nicks, since the DNA is kept in a double stranded conformation. However, during sequencing, single strand DNA is threaded through the pores and nucleotides after a nick are lost. A DNA repair step during library preparation did not prevent a shorter mean read length for this sample, hence use of fresh DNA is recommended. Nevertheless, our method of squiggle-based VNTR length estimation was robust to shorter read lengths.

### Tandem repeat characterization

Expanded alleles of the *ABCA7* VNTR have a strong risk increasing effect on AD, but characterization is restricted by the limitations of Southern blotting^22^. Here, we aimed to provide a better alternative based on PromethION sequencing. We first evaluated the classical sequencing analysis paradigm: base calling of raw sequencing data followed by alignment to a reference genome and ultimately (tandem repeat) variant calling. Base calling of raw ONT sequencing data is based on recurrent neural networks. To obtain very high accuracy, these machine learning algorithms require comprehensive training datasets which should ideally account for all possible sequence compositions and nucleotide modifications affecting current measurements. Unfortunately, this stage has not been reached, resulting in reduced base calling accuracy^15^. Several approaches exist to improve accuracy such as Nanopolish^44^, however all of these require the aid of a reference genome, which is not available for (expanded) tandem repeats. The lack of representative reference genomes also hinders reliable alignment of reads with tandem repeat sequences. As a result, characterization of the *ABCA7* VNTR using this canonical approach was unsatisfactory. Concerning TR length, we observed low accuracy and precision for *Albacore* and *Scrappie events* base callers. *Scrappie raw* obtained better results in this context, but failed to traverse most expanded VNTR alleles, which are crucial from a clinical point of view. On a nucleotide level, the erroneous base calls precluded reliable detection of alternative TR unit motifs. Particularly *Albacore* produced a deviating sequence composition, which has been observed in other GC-rich TRs^21^.

To overcome these issues, we developed NanoSatellite, a DTW based algorithm to perform TR variant calling directly on the raw squiggle level. We observed a strong correlation between Southern blotting lengths and repeat lengths estimated from the PromethION sequencing reads. The small differences in length estimation between both technologies can in part be explained by the limited resolution of Southern blotting. In addition, squiggle-based estimates had high precision, and performed well across all sequencing lengths, particularly expanded VNTRs. Moreover, NanoSatellite allowed investigation of TR nucleotide composition through direct comparison of raw TR unit squiggles instead of derivative base calls. We were able to observe clear differences in squiggle signals that could be attributed to nucleotide substitutions and insertions. Sequencing reads originating from the same VNTR allele showed highly consistent patterns of (alternative) TR unit squiggle motifs, hence validating that this squiggle-based method can be used to determine the sequence of TRs. Based on these sequences, we observed two VNTR alleles for individuals that appeared to have only one fragment on Southern blotting, thereby confirming that we spanned all VNTR alleles in all sequenced individuals.

NanoSatellite requires only genome coordinates of a TR of interest to generate reference squiggles. The TR delineation, segmentation, clustering, and sequence reconstruction are executed in an unsupervised fashion with a minimum of arbitrary cutoffs. To achieve the highest accuracy we limit analysis to biclustering on the strongest squiggle differences. Further sub-clustering to identify more TR motif interruptions and potentially nucleotide modifications is possible, yet requires supervised decisions and validation to balance the number of detected variants and accuracy. To facilitate further development, the algorithm is freely available and written in the statistical programming language R, which provides strong support of DTW functionality. To speed up computation, however, eventually switching to a lower-level programming language would be preferred. Nevertheless, we achieved high single-read accuracy for both TR length and sequence, which opens novel avenues in TR research. Many TRs in the human genome - some of which are currently uncharacterized - can be studied at once with a single sequencing run and somatic differences of unstable (expanded) TRs could be evaluated, which eventually will lead to the identification of novel disease-associated TRs and improved diagnostics.

## Acknowledgements

The authors thank the personnel of the Institute Born-Bunge, the neurological centers of the BELNEU consortium partners, and Oxford Nanopore Technologies, particularly Jonathan Pugh, James Platt, and Tim Massingham for their support on PromethION sequencing and base calling. The research was funded in part by the Alzheimer Research Foundation (SAO-FRA), the VIB Technology Fund, The Research Foundation Flanders (FWO), and the University of Antwerp Research Fund. ADR is recipient of a PhD fellowship of FWO, and a Hope in Head award from the Belgian Rotary. WDC is a recipient of a PhD fellowship from IWT/VLAIO.

## Authorship contributions

Conception & design: ADR, KS; sample acquisition: ADR, LB, RC, CVB; data generation: ADR, WDC, TDP, JVD, SDH, PDR, MS; data analysis: ADR, WDC; interpretation: ADR, CVB, KS; creation of software: ADR; drafting of the manuscript: ADR, KS; revision of the manuscript: WDC, LB, RC, TDP, JVD, SDH, PDR, MS, CVB

## Declarations

ADR and WDC have received travel reimbursement from Oxford Nanopore Technologies to speak at London Calling 2018 and ASHG 2018. Oxford Nanopore Technologies has provided consumables free of charge for sequencing of the NA19240 reference genome.

